# Nasal commensal, *Staphylococcus epidermidis* shapes the mucosal environment to prevent influenza virus invasion through Serpine1 induction

**DOI:** 10.1101/2020.08.03.235648

**Authors:** Ara Jo, Jina Won, Chan Hee Chil, Jae Young Choi, Kang-Mu Lee, Sang Sun Yoon, Hyun Jik Kim

## Abstract

Our recent study presented evidence that *Staphylococcus epidermidis* (*S. epidermidis*) was the most frequently encountered microbiome component in healthy human nasal mucus and that *S. epidermidis* could induce interferon (IFN)-dependent innate immunity to control acute viral lung infection. The serine protease inhibitor Serpine1 was identified to inhibit influenza A virus (IAV) spread by inhibiting glycoprotein cleavage, and the current study supports an additional mechanism of Serpine1 induction in the nasal mucosa, which can be regulated through *S. epidermidis* and IFN signaling. The exposure of *in vivo* mice to human *S. epidermidis* increased IFN-λ secretion in nasal mucosa and prevented an increase in the burden of IAV in the lung. *S. epidermidis*-inoculated mice exhibited the significant induction of Serpine1 *in vivo* in the nasal mucosa, and by targeting airway protease, *S. epidermidis-induced* Serpine1 inhibited the intracellular invasion of IAV to the nasal epithelium and led to restriction of IAV spreading to the lung. Furthermore, IFN-λ secretion was involved in the regulation of Serpine1 in *S. epidermidis*-inoculated nasal epithelial cells and *in vivo* nasal mucosa, and this was biologically relevant for the role of Serpine1 as an interferon-stimulated gene in the upper airway. Together, our findings reveal that human nasal commensal *S. epidermidis* manipulates the suppression of serine protease in *in vivo* nasal mucosa through Serpine1 induction and protects the nasal mucosa from IAV invasion through IFN-λ signaling.

**IMPORTANCE:** Previously, we proved that nasal microbiome could enhance IFN-related innate immune responses to protect the respiratory tract against influenza virus infection. The present study shows a great understanding of the intimate association of S. epidermidis-regulated IFN-lambda induction and serine protease inhibitor in nasal mucosa. Our data demonstrate that S. epidermidis-regulated Serpine1 suppresses the invasion of influenza virus through suppression of airway serine protease at the level of nasal mucosa and impedes IAV spread to the respiratory tract. Thus, human nasal commensal *S. epidermidis* represents a therapeutic potential for treating respiratory viral infections via the change of cellular environment in respiratory tract.

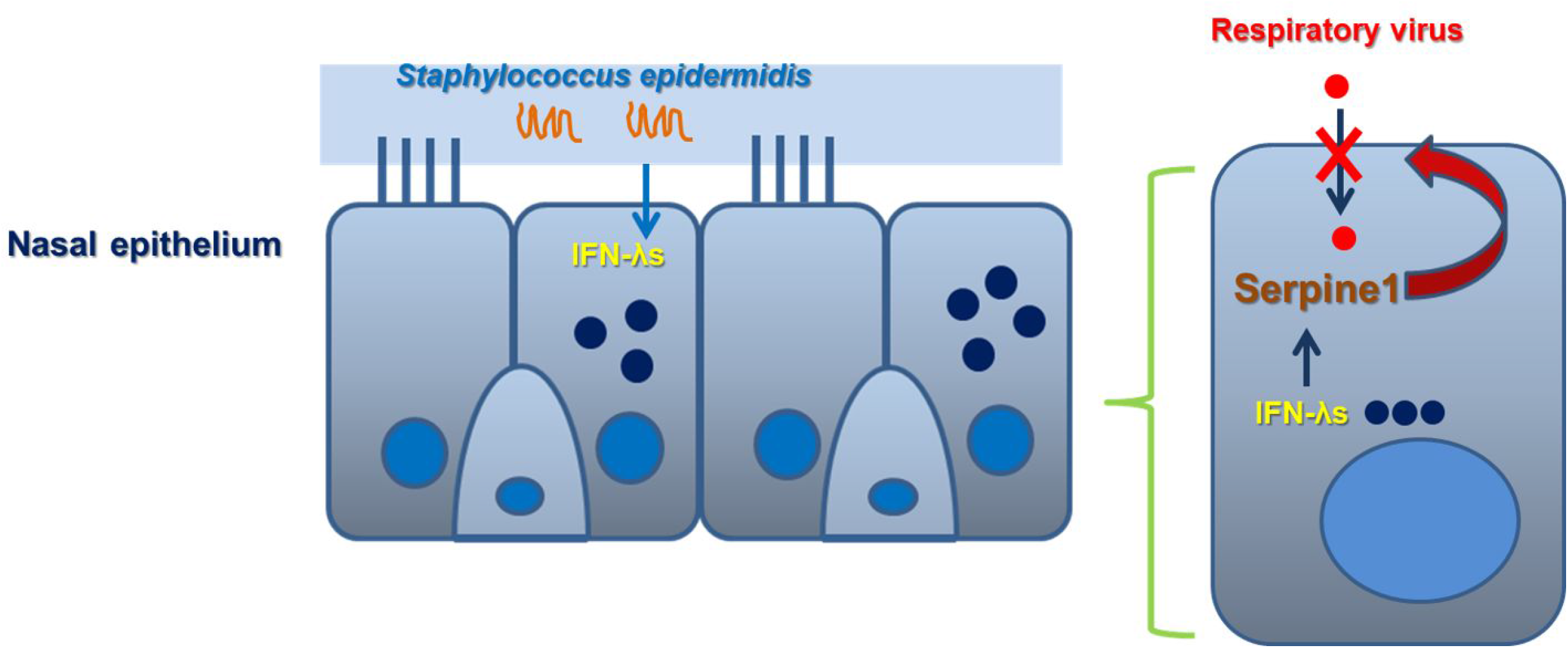

## Introduction

It is becoming increasingly apparent that the primary target cells of influenza A virus (IAV) infection are the epithelial cells of the respiratory tract, and the innate immune system of the respiratory epithelium serves as a first line of defense against invading respiratory viruses (1–3). It senses microbial molecules, such as single- and doublestranded viral RNA, and initiates the production of antiviral mediators such as interferon (IFN). IFNs are defined by their ability to induce resistance to viral infection in the respiratory tract (4–5). Type I and type III IFNs are directly produced in response to viral infection and contribute to the clearance of viral infections in the respiratory tract (7–9). Recently, it has been verified that type III IFNs (IFN-λ1, λ2, and λ3) are primarily responsible for protection against viral invaders in the respiratory epithelium and play an important role in local antiviral innate immunity (10–11).

Human mucosal surfaces are in direct contact with the external environment and are, therefore, susceptible to invasion and colonization by various pathogens (8). Studies on the clear reaction of the mucosal microbiome with the host increasingly take into consideration the contribution of mucosal immune responses and specific microbiome-mediated defense mechanisms against external pathogens (12). Respiratory mucosa, including those of nasal passages, are constantly exposed to inhaled pathogens, which directly impact the mucosal immune mechanisms (9,10). Inhaled pathogens encounter the host immune system for the first time in the respiratory mucosa, especially the nasal passage, and the microbial characteristics of the nasal mucus directly impact the mechanisms of initial immune responses (12, 13). Thus, insights into the microbiota of the human nasal mucosa can provide fundamental information regarding susceptibility to respiratory viral infections and factors contributing to related immune mechanisms, such as the induction of IFNs (14–16).

Our previous study identified *Staphylococcus epidermidis* as its most abundant constituent and showed that *S. epidermidis* that were isolated from healthy human nasal mucus accelerated the clearance of IAV from nasal epithelium through IFN-λ-related immune responses. Furthermore, human nasal commensal *S. epidermidis* prevents IAV lung infection in mice by enhancing IFN-λ-related innate immune responses in the nasal mucosa (17).

Many respiratory virus families require a maturation cleavage of viral surface glycoproteins, generally realized by serine proteases and a direct function of serine proteases in the respiratory tract, which induces the maturation of influenza virus to enhance cellular infectivity to the respiratory epithelium (18). It has been previously described that a serine protease inhibitor suppresses influenza virus maturation for intracellular invasion by targeting host proteases needed for viral glycoprotein cleavage (19–22). In this regard, it will be of immediate interest to determine whether serine protease inhibitor deficiency might be linked with the aggravation of human influenza virus replication and whether the localized administration of serine protease inhibitor to the respiratory tract might provide a new therapeutic approach for treating influenza viruses that require extracellular protease-driven maturation (22). We have already presented evidence of a key mechanistic link between the susceptibility to viral infections and nasal microbiome-mediated innate immunity. Here, we describe the more definite capacity of *S. epidermidis* in the human nasal microbiome to potentiate antiviral innate immune responses in the nasal epithelium through the induction of a serine protease inhibitor. These data indicate that *S. epidermidis* creates an intracellular environment in the nasal epithelium that makes it difficult for IAV to spread to the respiratory tract and can be a biological antiviral arsenal against invading influenza virus through the induction of Serpine1 driven by IFN-λ secretion.

## Results

### Nasal microbiome might be critical for suppression of IAV-caused lung infection

We first assessed whether the nasal microbiome could be important for its anti-viral protective properties using a murine model of IAV-caused lung infection. The nasal cavities of C57BL/6 (B6) mice (N=5) were inoculated with 30 μl of an antibiotic cocktail (vancomycin, neomycin, ampicillin, and metronidazole) (days 1, 2, and 3) following IAV infection through intranasal delivery (day 5); we then compared their lung pathologic findings with those of mice infected with IAV without intranasal antibiotics inoculation (Fig. 1A).

**FIG 1.**
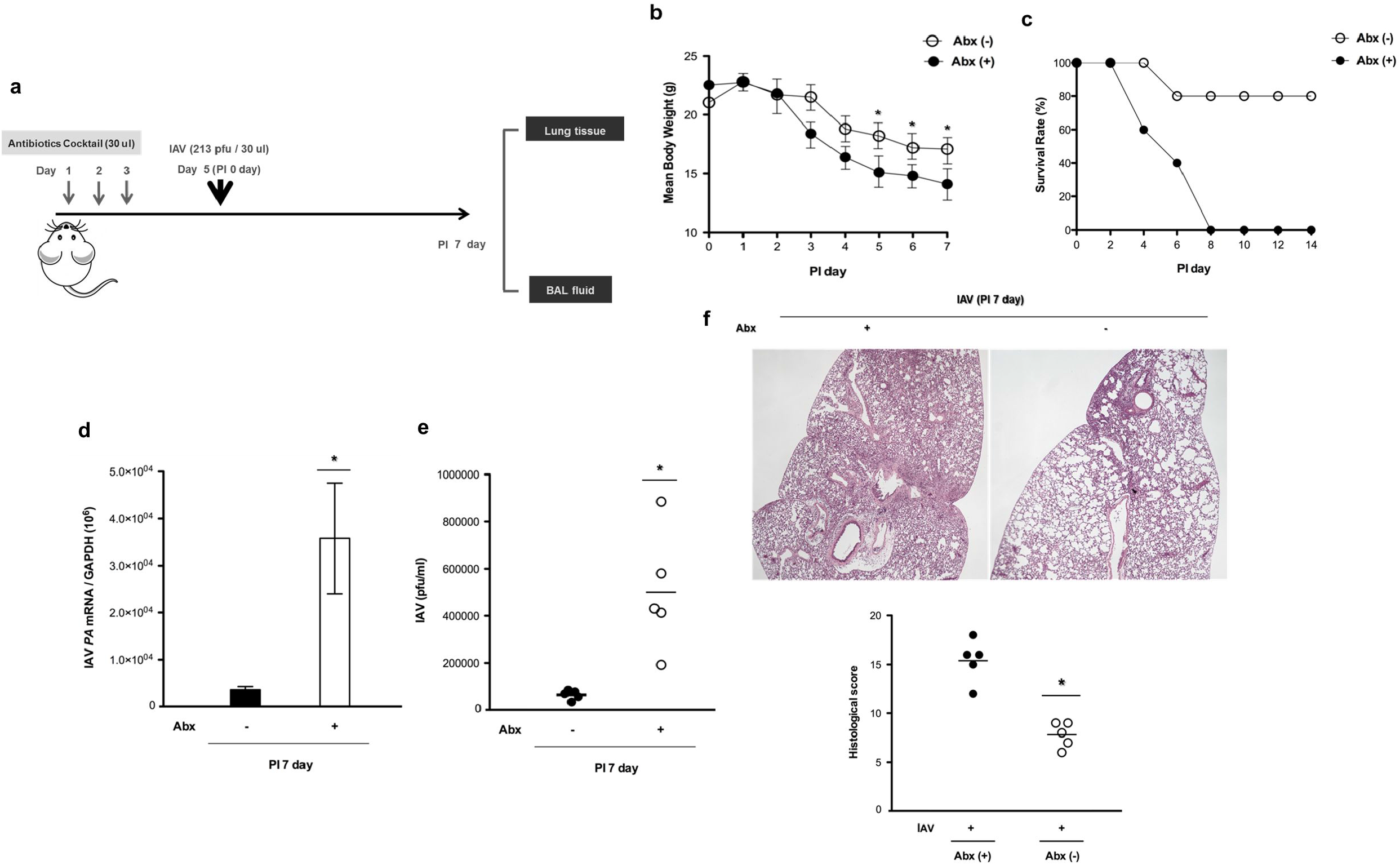
Nasal microbiome might be critical for suppression of IAV-caused lung infection. (a) Schematic of the mouse model experimental design with depletion of nasal microbiome. The native microbiome of the C57BL/6J mice was depleted with an antibiotic regimen (vancomycin, neomycin, ampicillin, and metronidazole) at 2 days prior to IAV infection. The mice (N=5) were infected with IAV (213 pfu) at the indicated time points. (b) Mean body weight and (c) survival rate of IAV-infected mice with or without intranasal antibiotics inoculation were measured. (d) IAV *PA* mRNA levels in the mice lung tissue and (e) viral titers from the BAL fluid of IAV-infected mice were assessed at 7 days post-infection (dpi). (f) Hematoxylin and eosin (H&E)-stained micrographs were also generated from lung sections obtained at 7 dpi. Micrographs shown are representative of lung sections from four mice. The micrographs were used to assess inflammation and tissue damage and to calculate a histological score. Real-time PCR and plaque assay results are presented as means ± SD from five independent experiments. \**p* < .05 compared to mice without antibiotics treatment. (Abx: antibiotics)

As gross determinants of virus-induced morbidity, the body weights and survival rates of the infected B6 mice were monitored for 14 days. The IAV-infected mice exhibited a significant decrease in mean body weight, with an 80% survival rate until 7 days post infection (dpi). Interestingly, the B6 mice that had antibiotics administered before the IAV infection did exhibit more significant weight loss than IAV-infected mice without antibiotics administration, and the mean body weight of that these mice lost (below 15 g until 7 dpi), resulted in the death of 100% of the mice after IAV infection (Fig. 1B and 1C).

Compared to the mice infected with IAV alone (3.2×10^3^), those inoculated with antibiotics preceding IAV infection showed higher IAV *PA* mRNA levels (3.4×10^4^) in the lung tissue (Fig. 1D) and higher viral titers (4.8×10^4^ [with antibiotics] vs 8.2×10^3^ [without antibiotics]) in BAL fluid (Fig. 1E). IAV-infected mice with antibiotics administration also showed more severe pathologic findings in the lung, with significantly higher histologic scores (Fig. 1F).

Together, these findings demonstrated that IAV-caused lung infection progressed more seriously in the case where the nasal microbiome was eliminated before infection and that the nasal microbiome can contribute to boosting the immune responses in the respiratory tract against IAV infection, thereby suppressing acute IAV lung infection.

### Pretreatment with human nasal microbiome *S. epidermidis* suppresses IAV-caused lung infection *in vivo*

Our previous study revealed that *S. epidermidis* is the most abundant microbiome component that colonizes healthy human nasal mucus and that the distribution of *S. epidermidis* might be significantly associated with innate immune responses in the nasal mucosa (17). Our findings also imply that the intranasal administration of *S. epidermidis* is a potential strategy for controlling viral replication at the level of the nasal mucosa. To clarify our hypothesis, we next studied whether the human nasal microbiome, *S. epidermidis* showed anti-IAV protective properties *in vivo* using a murine model of infection. The nasal cavities of B6 mice (N=5) were inoculated with human nasal mucus-derived *S. epidermidis* at 2 days (day 5) following nasal microbiota-depletion (day 1, 2, 3) using 30 μl of antibiotics. Then, the *S. epidermidis-inoculated* B6 mice were infected with IAV (213 pfu/30 μl) 2 days after *S. epidermidis* inoculation (day 7) (Fig. 2A).

**FIG 2.**
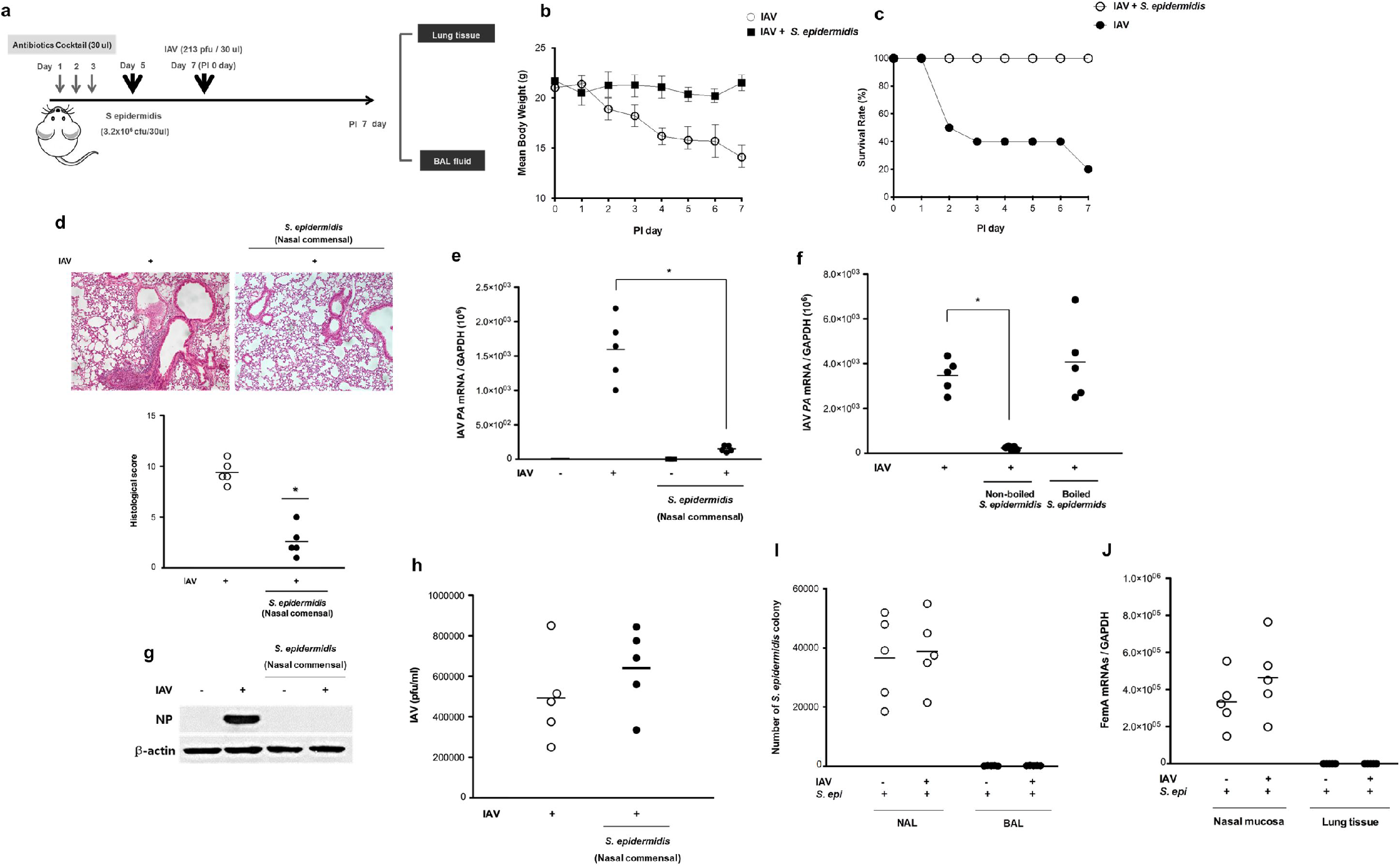
Human nasal mucus-derived *S. epidermidis* suppresses the viral spread and replication in IAV-infected mice. (a) Schematic of the mouse model experimental design for intranasal *S. epidermidis* inoculation and IAV infection. The native microbiome of the C57BL/6J mice was depleted with an antibiotic regimen prior to inoculation. The mice (N=5) were inoculated with *S. epidermidis* (3.2×10^6^ CFU/30 μl PBS) and/or with IAV (213 pfu) at the indicated time points. (b) Mean body weight and (c) survival rate of IAV-infected mice with or without *S. epidermidis* inoculation were measured. (d) B6 mice were inoculated with human nasal mucus-derived *S. epidermidis* prior to IAV infection, and H&E-stained micrographs were also generated from lung sections obtained at 7 dpi. Micrographs shown are representative of lung sections from five mice and were used to assess inflammation and tissue damage and to calculate a histological score. (e-f) B6 mice were inoculated with boiled and non-boiled human nasal *S. epidermidis* before IAV infection, and IAV *PA* mRNA levels in the mouse lung tissue were assessed at 7 dpi. (g) Levels of IAV NP were monitored in lung tissue using western blot analysis and representative results are shown. (h) Viral titers were also measured in the BAL fluid of IAV-infected mice following *S. epidermidis* inoculation. (i) *S. epidermidis* CFUs were determined at 7 dpi in the NAL and BAL fluid of IAV-infected mice (N=5) following *S. epidermidis* inoculation. (j) FemA mRNA of *S. epidermidis* were measured in the nasal mucosa and lung tissue of IAV-infected mice (N=5) at 7 dpi following *S. epidermidis* inoculation. Real-time PCR, plaque assays, and ELISA results are presented as means ± SD from five independent experiments. \**p* < .05 compared with mice infected with IAV alone.

As gross determinants of virus-induced morbidity, the body weights and survival rates of the infected B6 mice were monitored for 7 days. IAV-infected mice with intranasal antibiotics administration exhibited a significant decrease in mean body weight with a 20% survival rate until 7 dpi. Interestingly, the B6 mice that were administered human nasal mucus-derived *S. epidermidis* before IAV infection did maintain their body weight even after IAV infection, and the mean body weight of these mice exceeded 20 g until 7 dpi, resulting in 100% survival of the mice after the IAV infection (Fig. 2B and 2C). The significant pathologic findings were observed in the lung obtained from IAV-infected mice that were administered antibiotics before IAV infection. Compared to the IAV-infected mice with intranasal antibiotics administration, *S. epidermidis* exposure also resulted in attenuated pathologic findings in the lung of IAV-infected mice, with significantly lower histologic scores (Fig. 2D).

Human nasal microbiome *S. epidermidis* also showed potent antiviral activity by suppressing viral replication in the lungs of IAV-infected mice. Those infected with IAV following *S. epidermidis* inoculation showed lower IAV *PA* mRNA levels (2.7×10^2^) in the lung tissue of mice with antibiotics pretreatment (Fig. 2E) and IAV mRNA levels increased again in the lungs of IAV-infected mice that were treated with boiled *S. epidermidis* (4.02×10^3^) (Fig. 2F). The nucleoprotein level of IAV was significantly reduced in the lungs of IAV-infected mice that were inoculated with human nasal *S. epidermidis* compared to the IAV-infected mice pretreated with antibiotics (Fig. 2G). However, viral titers still remained significantly elevated in the BAL fluid of IAV-infected mice despite inoculation with human nasal *S. epidermidis* (Fig. 2H).

We next assessed the distribution of bacteria in *S. epidermidis*-inoculated mice by comparing the colony forming units (CFUs) in the NAL and BAL samples. Whereas substantial numbers of *S. epidermidis* CFUs were observed in the NAL samples, the levels of *S. epidermidis* cells in the BAL samples were under the detection limit (Fig. 2I). We also found that mRNA levels of S. *epidermidis* (FemA) were minimally detected in the lungs of mice after human nasal *S. epidermidis* inoculation (Fig. 2J).

These findings demonstrated that the human nasal microbiome *S. epidermidis* can boost an antiviral immune response at the level of mouse nasal mucosa, thereby suppressing IAV replication or spread to the respiratory tract and preventing acute IAV lung infection but viral titer in respiratory mucus of IAV-infected mice was not reduced by intranasal inoculation of *S. epidermidis.*

### Human nasal microbiome *S. epidermidis* promoted the induction of serine protease inhibitor in nasal epithelium

To analyze the effects of *S. epidermidis* inoculation on the susceptibility of the nasal mucosa to IAV infection, NHNE cells from five healthy subjects were inoculated with human nasal *S. epidermidis* at a multiplicity of infection (MOI) of 0.25. Subsequently, *S. epidermidis* mRNA levels in the cell lysate and the colony count of *S. epidermidis* in the supernatant were assessed in *S. epidermidis-inoculated* NHNE cells at different time points post-infection. Real-time PCR revealed that *S. epidermidis FemA* mRNA levels increased significantly starting from 8 hr post infection and that the highest levels were observed at 1 dpi (8 hr: 8.2×10^4^;1 dpi: 5.8×10^5^; Fig. 3A). The numbers of *S. epidermidis* CFUs were also significantly increased in the supernatant of *S. epidermidis-inoculated* NHNE cells until 1 day after *S. epidermidis* inoculation (Fig. 3B).

**FIG 3.**
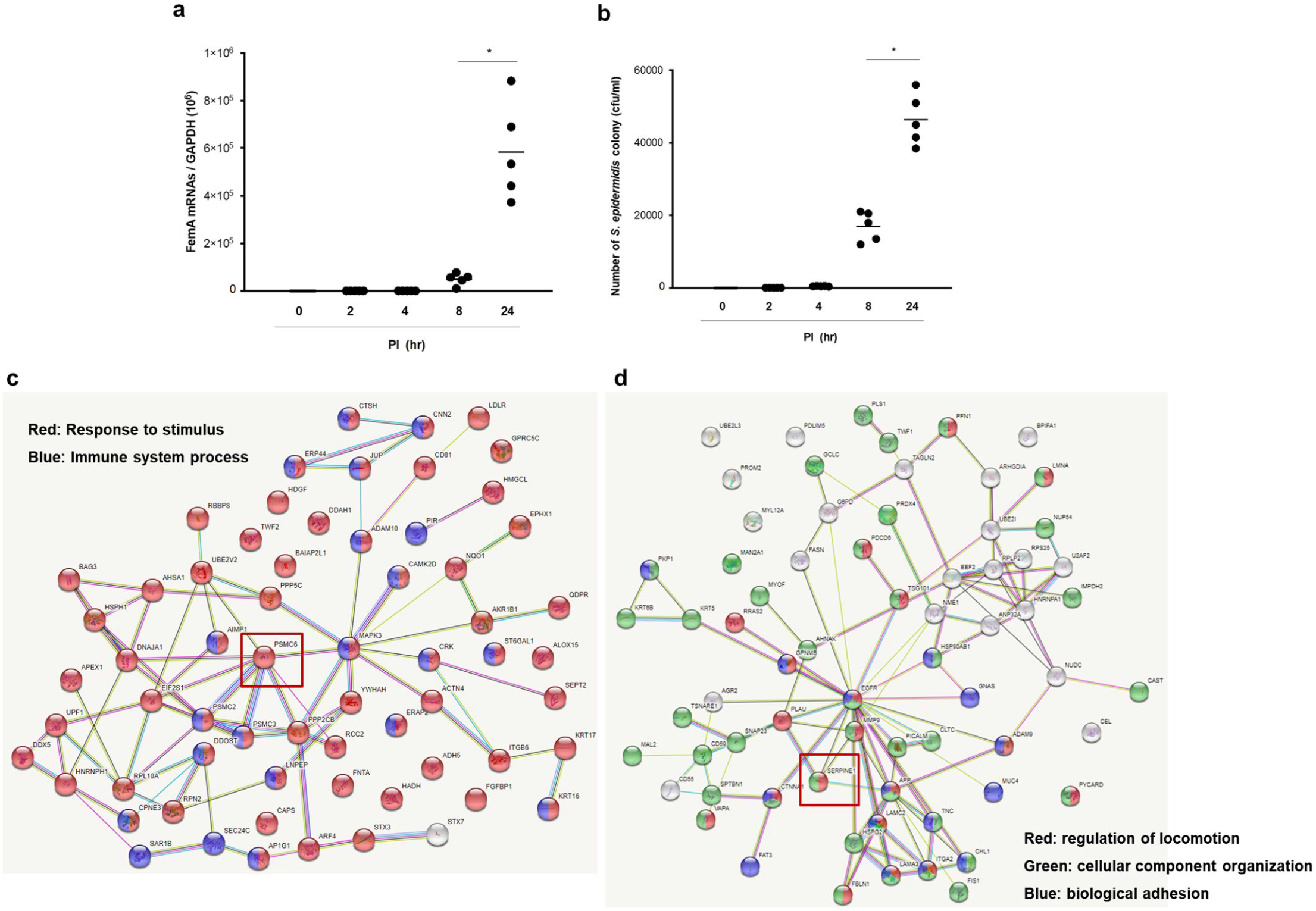
Human nasal microbiome *S. epidermidis* promoted the induction of serine protease inhibitor in the nasal epithelium. NHNE cells from five healthy volunteers were inoculated with human nasal *S. epidermidis* at a MOI of 0.25. (a) FemA mRNA levels, normalized to cellular *GAPDH* transcript levels, were monitored by real-time PCR over the course of 3 days. (b) *S. epidermidis* CFUs were determined at 1 dpi in the supernatant of S. epidermidis-inoculated NHNE cells. Proteomics analysis through LC-MS/MS analyses with the aid of a Quadrupole Orbitrap mass spectrometer and STING analysis were performed to evaluate the changes in (c) the proteins which were identified only in the supernatant of *S. epidermidis*-inoculated NHNE cells and (d) the proteins which were detected over 4 times in the supernatant of *S. epidermidis*-inoculated NHNE cells). Results are presented as the means ± SD from five independent experiments. \**p* < .05 compared with mock-infected NHNE cells.

To evaluate the changes in protein expressions in the nasal epithelium after *S. epidermidis* inoculation, we performed proteomics analyses through LC-MS/MS analyses (data-dependent acquisition [DDA]) with the aid of a Quadrupole Orbitrap mass spectrometer (Q-Exactive Plus; Thermo Fisher Scientific, Waltham, MA, USA). The supernatant was harvested from NHNE cells and *S. epidermidis*-inoculated NHNE cells for 24 hr from 5 independent cultures. The mean protein concentration of the supernatant of NHNE cells was 2.56 + 1.95 mg/ml, and it was 0.95 + 0.84 mg/ml in the supernatant of *S. epidermidis*-inoculated NHNE cells. After filtering out contaminants and proteins without triplicate intensity values in the supernatant of NHNE with and without *S. epidermidis* inoculation, in total, 939 unique proteins were identified in both supernatants, corresponding to 821 proteins in the supernatant of the NHNE cells without *S. epidermidis* inoculation (control), and 886 proteins in the supernatant of *S. epidermidis*-inoculated NHNE cells, and they were retained for differential abundance analysis (Fig. S1). A total of 768 proteins were classified in the supernatants from both control and *S. epidermidis*-inoculated NHNE cells, and 496 proteins were found to be differentially abundant. A 2-sample *t*-test with a fold change cutoff at approximately 4 was used to highlight proteins that characterized the control NHNE cells and *S. epidermidis-inoculated* NHNE cells. Among them, 122 proteins were more than 4-fold higher in expression than control NHNE cells and found to be differentially abundant. A total of 118 proteins were highlighted into only the supernatant of the *S. epidermidis*-inoculated NHNE cells, and 114 proteins were found to be differentially abundant. To observe the distribution of differentially abundant proteins in the supernatant of NHNE cells depending on *S. epidermidis* inoculation, we used gene ontology analysis (biological process by DAVID). The analyzed proteins were classified into 2 groups: proteins that were detected only in the supernatant of *S. epidermidis-inoculated* NHNE cells and proteins which were detected at levels over 4-fold higher in the supernatant of *S. epidermidis-inoculated* NHNE cells compared to the supernatant of control NHNE cells (Fig. S2 and S3).

Considering that the IAV mRNA level was decreased but the viral titer was not reduced after *S. epidermidis* inoculation, we thought that *S. epidermidis* disturbed the invasion of IAV to the respiratory mucosa and focused on the induction of serine protease inhibitor in the *S. epidermidis*-inoculated NHNE cells resulting in suppression of serine protease-regulated viral maturation. STRING database analysis revealed that there were two proteins that were related to serine protease inhibitor in both supernatants. *Homo sapiens* serpin peptidase inhibitor (PSMC6) was identified in the supernatant of *S. epidermidis*-inoculated NHNE cells including the response to stimulus GO category in group 1 (Fig. 3C) and plasminogen activator inhibitor 1 (Serpine1) in group 2 was 11.07-fold higher than that in the supernatant of NHNE cells without *S. epidermidis* inoculation including the regulation of locomotion and cellular component organization GO category (Fig. 3D).

These results showed that the secreted protein levels of serine protease inhibitor were induced in human nasal microbiome *S. epidermidis-inoculated* nasal epithelium and that *S. epidermidis* might inhibit the cellular invasion of IAV, resulting in decreased viral spreading to the respiratory tract.

### Serpine1 was induced dominantly in *S. epidermidis*-inoculated nasal epithelium and suppressed the activity of intracellular serine protease

We studied the kinetics of expression of human *Serpine* genes such as *Serpine1, Serpine2,* and *Serpine3,* as well as protein production and secretion in NHNE cells after inoculation with human nasal microbiome *S. epidermidis* (MOI 0.25). The PCR results showed that *Serpine1* mRNA was significantly upregulated 8 hr after *S. epidermidis* inoculation (4.8×10^4^) compared to NHNE cells without S. epidermidis inoculation (1.2×10^3^) (Fig. 4A). Both *Serpine2* and *Serpine3* gene expression levels were minimally induced in NHNE cells after *S. epidermidis* inoculation. Consistent with the induction of *Serpine1* gene expression, increased intracellular and secreted protein levels of Serpine1 were significantly elevated in *S. epidermidis*-inoculated NHNE cells, and their highest levels were observed at 24 hr after inoculation in western blot analysis and ELISA results (Fig. 4B and 4C). To assess the effects of *S. epidermidis*-induced Serpine1 expression on the susceptibility of the nasal epithelium to IAV infection, NHNE cells were inoculated with *S. epidermidis* at MOI 0.25 and then infected with IAV 8 hr after inoculation at MOI 1. Interestingly, PCR results revealed that *Serpine1* mRNA was also upregulated in IAV-infected NHNE cells (1.2×10^4^ + 4.3×10^3^) at 1 dpi and *Serpine1* gene expression was also induced by *S. epidermidis* inoculation (1.1×10^4^ + 3.1×10^3^). However, *Serpine1* gene expression level was significantly higher in NHNE cells with *S. epidermidis* inoculation following IAV infection (3.9×10^4^ + 5.2×10^3^, Fig. 4D). Subsequently, the IAV viral titer was assessed in the supernatants of *S. epidermidis*-inoculated NHNE cells following IAV infection depending on the neutralization of Serpine1 in NHNE cells. The plaque assay revealed that increased viral titer (5.2×10^7^ pfu/ml) after IAV infection was completely reduced in the supernatant of IAV-infected NHNE cells with *S. epidermidis* inoculation (8.1×10^4^ pfu/ml), but the supernatant of IAV-infected NHNE cells treated with Serpine1 neutralizing antibody exhibited increased viral titer (2.4×10^7^ pfu/ml) again regardless of *S. epidermidis* inoculation (Fig. 4E). Western blot analysis similarly revealed that IAV NP level was not reduced in IAV-infected NHNE cells that were treated with Serpine1 neutralizing antibody before *S. epidermidis* inoculation (Fig. 4F).

**FIG 4.**
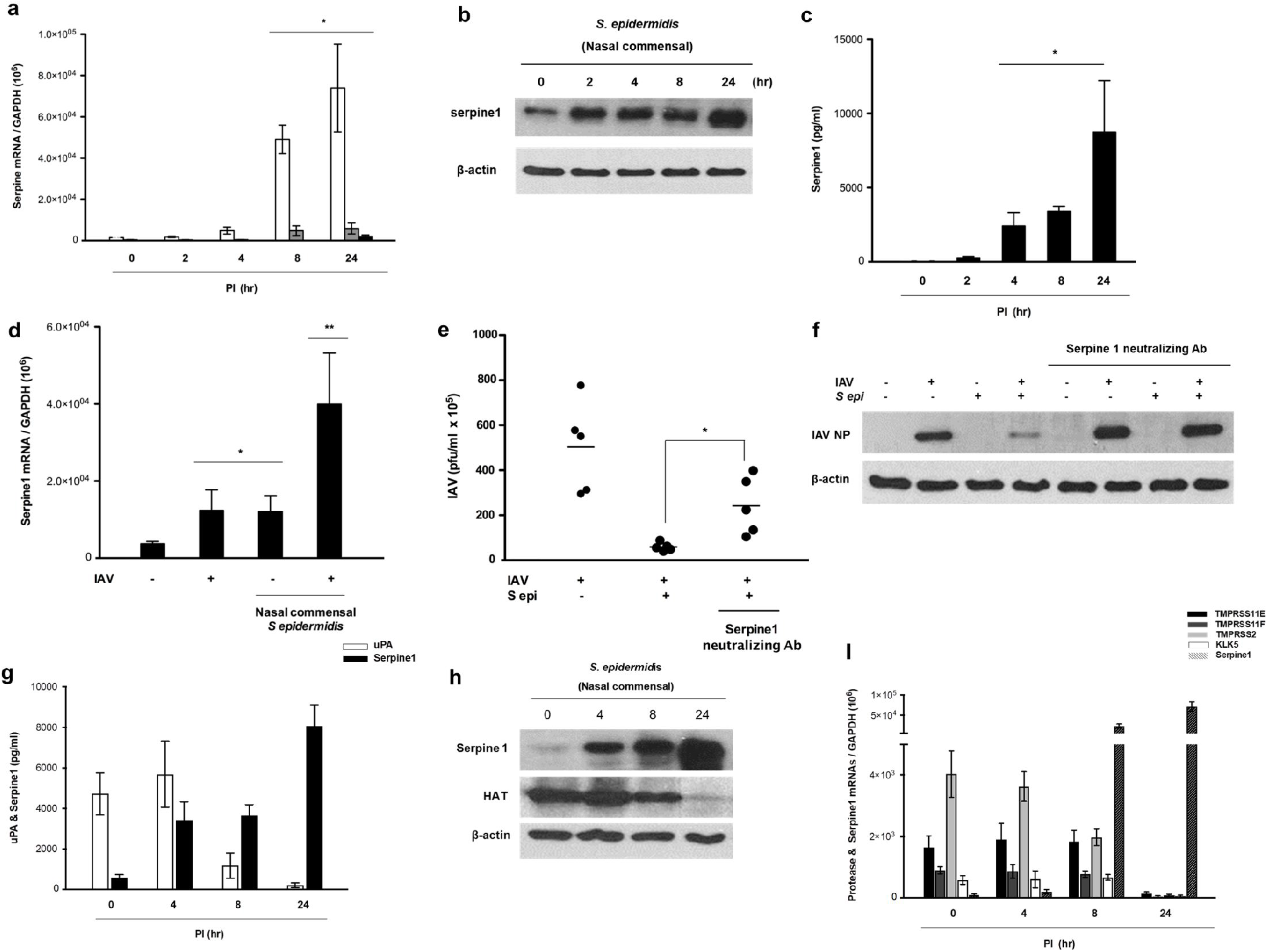
Serpine1 was induced dominantly in *S. epidermidis-inoculated* nasal epithelium and suppressed the activity of intracellular serine protease. NHNE cells from five healthy volunteers were inoculated with human nasal *S. epidermidis* at a MOI of 0.25. (a) Serpine1, Serpine2, and Serpine3 mRNA levels, normalized to cellular *GAPDH* transcript levels, were monitored by real-time PCR over the course of 1 day. (b) The intracellular protein level of Serpine1 was measured in the cell lysate of *S. epidermidis*-inoculated NHNE cells at 1 dpi using western blot analysis. (c) The secreted protein level of Serpine1 was measured in the supernatant of *S. epidermidis*-inoculated NHNE cells at 1 dpi using ELISA. (d) Serpine1 mRNA level, normalized to cellular *GAPDH* transcript levels, was monitored by real-time PCR over the course of 1 day after IAV infection following *S. epidermidis* inoculation. (e-f) The neutralizing antibody for Serpine1 was administered to NHNE cells 1 hr prior to *S. epidermidis* inoculation, and then the cells were infected with IAV for 1 day. Viral titers and NPs of IAV were compared in IAV-infected NHNE cells after *S. epidermidis* inoculation depending on the neutralization of Serpine1. (g-h) The intracellular protein levels of uPA, HAT, and Serpine1 were measured in the cell lysate of *S. epidermidis*-inoculated NHNE cells at 4, 8, and 24 hr using ELISA and western blot analysis. (i) The mRNA levels of proteases such as TMPRSS2, TMPRSS11E, TMPRSS11F, and KLK5, which are targeted by Serpine1, were measured using the cell lysate of *S. epidermidis*-inoculated NHNE cells at 4, 8, and 24 hr. The figure of the western blot is representative from five independent experiments. Real-time PCR and plaque assay results are presented as means ± SD from five independent experiments. \**p* < .05 compared with mock-infected NHNE cells.

Infectivity of IAV progeny particles requires a maturation cleavage of the viral hemagglutinin, catalyzed by host proteases (22). We tested whether *S. epidermidis*-induced Serpine1 might target for host protease reduction in NHNE cells and inhibit the maturation of IAV in the nasal epithelium.

First, the secreted protein level of urokinase plasminogen activator (uPA), a major target of Serpine1 protein, was measured, and ELISA results showed that the concentration of secreted uPA protein was gradually reduced in the supernatant of *S. epidermidis*-inoculated NHNE cells until 24 hr after inoculation. In contrast, the concentration of Serpine1 protein was significantly increased after *S. epidermidis* inoculation, showing the highest level at 24 hr after inoculation (Fig. 4G). Furthermore, human airway trypsin (HAT) has been also identified as an established target of Serpine1 protein, and western blot analysis showed that HAT expression decreased significantly in the cell lysate of NHNE cells after *S. epidermidis* inoculation. In contrast, we were able to detect the gradual increase of Serpine1 protein expression, and its highest expression was observed at 24 hr after *S. epidermidis* inoculation (Fig. 4H). We also found similar results for the gene expression of extracellular airway proteases such as transmembrane protease serine (TMPRSS)2, TMPRSS11E, TMPRSS11F, and KLK5, and the real-time PCR data likewise revealed that gene expression levels of extracellular airway proteases were gradually reduced in NHNE cells after *S. epidermidis* inoculation, in contrast to the gene expression of *Serpine1* (Fig. 4I). These findings suggested that nasal commensal *S. epidermidis* induces Serpine1 production and secretion and targets the reduction of airway proteases needed for viral maturation in the nasal epithelium.

### IFN-λ contributes to *S. epidermidis-induced* Serpine1 production in nasal epithelium

Previously, we demonstrate that the levels of *S. epidermidis* in human nasal mucus proportionally affect the transient expression of IFN-λ in nasal mucosa and human nasal commensal *S. epidermidis* can boost the IFN-λ-dependent innate immune response in mouse nasal mucosa, thereby suppressing IAV replication at the level of the nose and preventing acute IAV lung infection (17).

To assess the mechanisms of the *S. epidermidis-dependent* Serpine1 production upon IFN-λ stimulation in the nasal epithelium, we measured the expression of Serpine1 in *S. epidermidis*-exposed NHNE cells depending on IFN-λ treatment. First, NHNE cells were treated with recombinant IFN-λ (IFN-λ1: 10 ng/ml and IFN-λ2: 10ng/ml) for 2, 4, 8, and 24 hr, and both *Serpine1* mRNA and Serpine1 protein levels were measured in the cell lysate and supernatant of the IFN-λ-inoculated NHNE cells. The real-time PCR results showed that *Serpine1* mRNA was elevated starting 8 hr (3.8 × 10^4^ + 1.8 × 10^3^) after IFN-λ inoculation (Fig. 5A), and ELISA data revealed that the secreted protein of Serpine1 was also increased until 24 hr in IFN-λ-inoculated NHNE cells (Fig. 5B). In addition, the intracellular protein expression of Serpine1 was significantly induced in the IFN-λ-inoculated NHNE cells, and the highest protein level was observed at 24 hr after inoculation (Fig. 5C).

**FIG 5.**
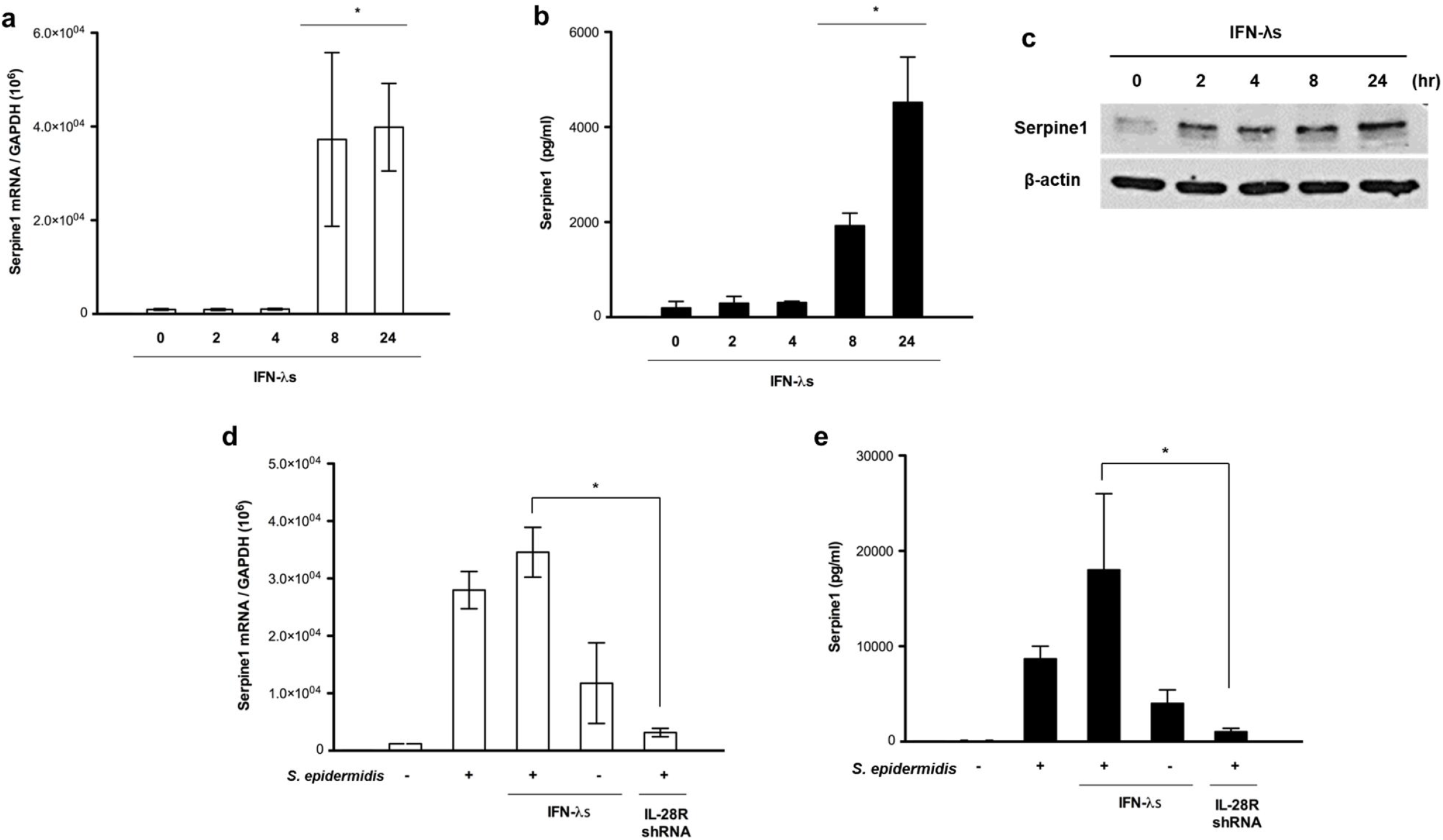
IFN-λ contributes to *S. epidermidis-induced* Serpine1 production in the nasal epithelium. (a-c) NHNE cells were treated with recombinant IFN-λ (IFN-λ1:10 ng/ml and IFN-λ2: 10 ng/ml) for 24 hr, and *Serpine1* mRNA, secreted protein, and intracellular protein levels were monitored by real-time PCR, ELISA, and western blot analysis. (d-e) *Serpine1* mRNA and secreted protein levels were also measured in NHNE cells that were transfected with control shRNA or IL28R shRNA. The figure of a western blot is representative from five independent experiments. PCR and ELISA results are presented as the means ± SD from five independent experiments. \**p* < .05 compared with control NHNE cells.

As a next step, NHNE cells were transfected with short hairpin (sh)RNA of IL28R, which is a region of the IFN-λ receptor, to cause a functional loss of the IFN-λ-related signaling pathway. Interestingly, the increase in the *Serpine1* mRNA level in *S epidermidis*-inoculated NHNE cells (3.8×10^4^) was significantly attenuated in NHNE cells with transfection of *IL28R* shRNA (3.2×10^3^) before *S. epidermidis* inoculation (Fig. 5D). ELISA results showed that *S. epidermidis*-induced secreted protein levels of Serpine1 were also reduced in the supernatant of NHNE cells with transfection of *IL28R* shRNA (Fig. 5E).

These data suggested that nasal commensal *S. epidermidis* induces Serpine1 production in the nasal epithelium and that IFN-λ promotes *S. epidermidis*-regulated Serpine1 transcription and secretion.

### *S. epidermidis* induces Serpine1 production *in vivo* in nasal mucosa

To investigate further the role of Serpine1 in the nasal mucosa of IAV-infected mice, B6 mice (N=5) were inoculated with human nasal mucus-derived *S. epidermidis* through the nasal cavity at 2 days (day 5) following nasal microbiota-depletion (day 1, 2, 3) using 30 μl of antibiotics. Then, *S. epidermidis*-inoculated B6 mice were infected with IAV (213 pfu/30 μl) at 2 days after *S. epidermidis* inoculation (day 7). Human nasal microbiome *S. epidermidis* also showed antiviral potential by suppressing viral replication in the nasal mucosa of IAV-infected mice. Those infected with IAV following *S. epidermidis* inoculation showed lower IAV *PA* mRNA levels (0.4×10^3^) in the lung tissue (Fig. 6A), and IAV NP levels significantly decreased in the nasal mucosa of IAV-infected mice that were treated with *S. epidermidis* compared to IAV-infected mice with antibiotics pretreatment (Fig. 6B). However, viral titers still remained higher in the NAL fluid of IAV-infected mice despite inoculation with human nasal *S. epidermidis* (Fig. 6C). These findings suggested that the presence of *S. epidermidis* might suppress the invasion of IAV *in vivo* in the nasal mucosa with the decrease of intracellular IAV expression and maintenance viral titer in NAL fluid. Actually, *Serpine1* was transcriptionally upregulated in the nasal mucosa of B6 mice upon IAV stimulation, and *Serpine1* gene expression was higher in the nasal mucosa of B6 mice with inoculation of *S. epidermidis* (Fig. 6D). In addition, the secreted protein level of Serpine1 was also enhanced depending on the presence of *S. epidermidis* in the nasal mucosa of B6 mice (Fig. 6E). These results strengthen the physiological relevance of Serpine1 as regulator of IAV invasion in nasal mucosa of IAV-infected mice and a component of the host defense factor that restricts IAV spread to respiratory tract from upper airway of *in vivo.*

**FIG 6.**
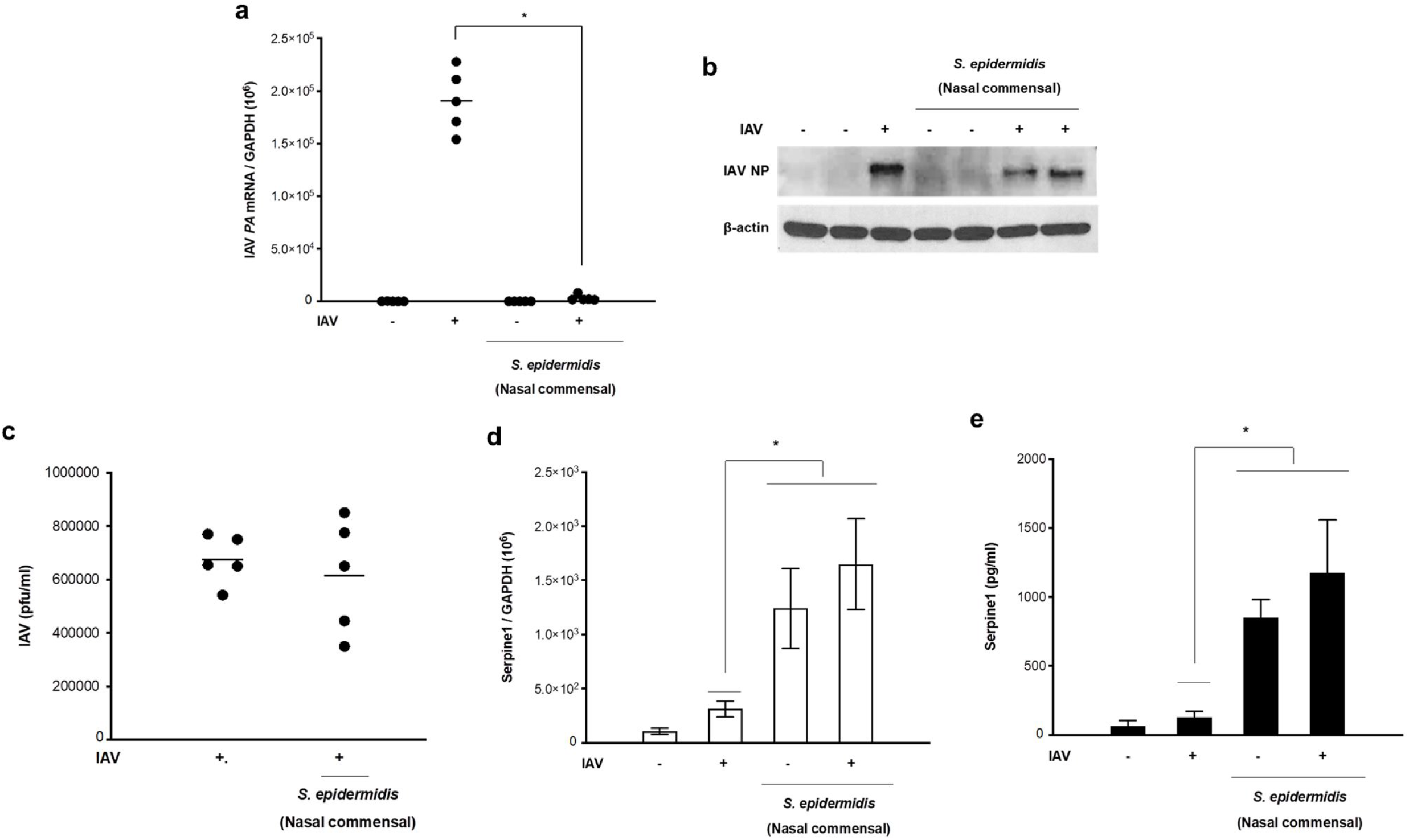
*S. epidermidis* induces Serpine1 production *in vivo* in the nasal mucosa. The native microbiome of the C57BL/6J mice was depleted with an antibiotic regimen prior to inoculation. The mice (N=5) were then inoculated with *S. epidermidis* (3.2×10^6^ CFU/30 μl PBS) and/or with IAV (213 pfu) at the indicated time points. (a-b) IAV *PA* mRNA and IAV NP levels in the mouse nasal mucosa were assessed at 7 dpi using real-time PCR and western blot analysis. (c) Viral titers were also measured in the NAL fluid of IAV-infected mice following *S. epidermidis* inoculation. (d) Serpine1 mRNA and (e) secreted protein levels were also monitored in the mouse nasal mucosa and NAL fluid using real-time PCR and ELISA at 7 dpi. The figure of a western blot is representative from five independent experiments. PCR and ELISA results are presented as the means ± SD from five independent experiments. \**p* < .05 compared with the results from control B6 mice.

## Discussion

Our study revealed that *S. epidermidis,* which colonizes abundantly healthy human nasal mucus, might be significantly associated with the induction of serine protease inhibitor, Serpine1 in nasal mucosa and could restrict IAV spread to the lung by targeting serine proteases for IAV maturation in the upper airway. Our findings also imply that Serpine1 is identified as an IFN-stimulated gene for the prevention of respiratory viral infections by disturbing viral maturation and that IFN-λ could mediate *S. epidermidis*-regulated Serpine1 induction at the level of the nasal mucosa. This study presents novel evidence of the role played by the healthy human nasal microbiome in the host defense mechanism against IAV spread to the respiratory tract.

Respiratory mucosa is the first target organ for environmental pathogens, including respiratory viruses and recent works have highlighted the critical role of the respiratory mucosa as a barrier for restricting the invasion of the host by multiple pathogens (10, 23,24). The compositional and predicted functional differences in the respiratory microbiome resulting from environmental stimuli have been drawing increasing interest, and the importance of the respiratory microbiome, especially with respect to immune protection, has been significantly recognized (25–30). The nasal mucosa is also a key player in immunological defense to protect the respiratory tract and is responsible for the filtration of inhaled pathogens from direct exposure to pressurized airflow (24,31). Human respiratory viruses first encounter host defense mechanisms in the nasal epithelium, and host protection against viral infections can be conferred by a specialized innate immune system of the nasal epithelium capable of combating invasion by respiratory viruses (10,32–37). There is also growing evidence that a microbiome community resides in human nasal mucus. Our previous study revealed that *S. epidermidis* is the most abundant commensal organism in the healthy human nasal mucus, and the presence of *S. epidermidis* contributes to strengthening the antiviral immune defense mechanisms of response in the respiratory tract (17). The current findings also demonstrated that IAV-caused lung infection progressed more seriously in the case where the nasal microbiome was eliminated before infection and that the nasal microbiome can contribute to boosting the immune responses in the respiratory tract against IAV infection, thereby suppressing acute IAV lung infection. However, the studies about more definite interplay between *S. epidermidis* and respiratory virus in nasal mucosa may be needed to prove the synergy of the mucosal immunity during the virus-caused respiratory infection

Antiviral innate immunity in the respiratory tract is mediated by an increase in IFN secretion (38–42), and IFN-regulated antiviral innate immune responses have been well-documented to activate the adaptive immune system (45–47). Increasing evidence shows that IFN-λ is also critical for antiviral innate immunity in the respiratory tract, with disrupted IFN-λ-related innate immunity increasing susceptibility to respiratory viral infections (48–51). We have also reported that IFN-λ represents the predominant IFN type induced by IAV and contributes to the first-line defense against viral infections in human nasal epithelial cells (10, 23, 52, 53). We have concentrated on verifying the immune factors mediated by IFN-λ in the nasal mucosa and have found that nasal microbiome *S. epidermidis* showed potent antiviral activity in the nasal epithelium and that all ensuing antiviral responses from *S. epidermidis* are dependent on the production of IFN-λ (17). Relative to infection with IAV alone, the inoculation of *S. epidermidis* before IAV infection reduced the viral burden of NHNE cells, while concomitantly inducing IFN-λ and ISG expression.

We demonstrate that the human nasal commensal *S. epidermidis* mediates front-line antiviral protection against IAV infection through modulation of IFN-λ-dependent innate immune mechanisms in the nasal mucosa, thereby demonstrating the role of host-bacterial commensalism in shaping human antiviral responses to impede the spread of respiratory virus. This symbiotic nasal microbiome, *S. epidermidis* creates an intracellular environment in which respiratory viruses are unlikely to survive and we understand that the difference of *S. epidermidis* abundance in nasal mucus explains the individual susceptibility to respiratory viral infection in each patient.

IAV maturation involves cleavages of its surface glycoprotein hemagglutinin (HA) into HA1 and HA2, and each HA subtype can be cleaved by a given protease that is determined by its cleavage site sequence. The glycoprotein cleavage of IAV is a major determinant of IAV pathogenicity, and the cleavage by extracellular airway proteases including serine protease is required for the infectivity or spread of IAV (54). Based on previous studies, IAV does not encode its own protease, and HA cleavage for IAV replication depends on the presence of host proteases at the respiratory tract. Therefore, HA cleavage has been proposed as a target for antiviral therapy, and the mediators that could induce anti-protease activity can be used for a new therapeutic option against IAV infection in the respiratory tract.

Our data showed that both IAV mRNA and NP levels were decreased in the nasal mucosa and lung of IAV-infected mouse after *S. epidermidis* inoculation to the nasal epithelium, but *S. epidermidis* inoculation did not reduce the viral titers of IAV in the BAL fluid and supernatant of nasal epithelium. Therefore, we thought that *S. epidermidis* disturbed the invasion of IAV to nasal epithelial cells, which resulted in a higher viral titer in the extracellular environment. Actually, we speculate that the nasal commensal *S. epidermidis* allowed it to boost the baseline mechanism for serine protease inhibitor through its own serine protease and inhibited IAV maturation through the induction of serine protease inhibitor in *S. epidermidis*-inoculated nasal epithelium.

The current proteomics data showed that the protein levels of the serine protease inhibitor Serpine1 and serpine peptidase inhibitor were significantly induced in the supernatant of *S. epidermidis-inoculated* NHNE cells. In addition, *in vitro* study revealed that human nasal microbiome *S. epidermidis* induced Serpine1 gene expression, intracellular protein, and secreted proteins in the cell lysate and supernatant of NHNE cells. Interestingly, *S, epidermidis*-induced Serpine1 expression in the nasal epithelium showed inverse correlation with the expression of airway proteases such as uPA, HAT, TMPRSS11E, TMPRSS11F, TMPRSS2, and KLK5, and the reduction of *S. epidermidis-induced* Serpine1 activity exhibited higher IAV replication in the nasal epithelium. These findings suggest that nasal microbiome *S. epidermidis* has a distinctive antiviral strategy against IAV through induction of serine protease inhibitor and shapes the nasal mucosa into an airway protease-deficient environment in the host. This mechanism of human nasal microbiome *S. epidermidis* functions to protect the respiratory tract of a host from natural IAV infection at the level of the nasal mucosa.

We presume that targeting serine protease shows the strong impact to reduce IAV spread from cell to cell, thus identifying Serpine1 as a candidate material that may influence on individual susceptibility to IAV and the outcome of infection. It is interesting that the upregulation of Serpine1 expression by *S. epidermidis* during IAV infection was induced depending on IFN-λ, and actually, we identified that Serpine1 might be an IFN-stimulated gene in the nasal epithelium. The current findings indicate that *S. epidermidis*, which predominantly resides in human nasal mucosa, induces IFN-λ expression as a host factor inhibiting IAV replication and that *S. epidermidis*-induced IFN-λ is primarily responsible for the production of Serpine1, resulting in mechanistically disturbing the maturation of IAV through the reduction of host airway proteases. In this regard, it will be of interest to determine whether induction of Serpine1 by nasal microbiome might be linked with the change of cellular environment to decrease IAV infectivity and the present study estimates that nasal microbiome *S. epidermidis*-regulated Serpine1 suppress the invasion of IAV to nasal epithelial cells, and thus may impede IAV replication in the human respiratory tract.

Our study provides a greater understanding of how the nasal microbiome enhances IFN-dependent innate immune responses to protect the respiratory tract against influenza viral infection. The abundant human nasal microbial organism *S. epidermidis* enhances resistance against IAV infection in human nasal epithelium and suppresses IAV invasion of the nasal epithelium through IFN-λ-dependent Serpine1 amplification. This intimate association of *S. epidermidis* with IFN-λ and serine protease inhibitor could potentially benefit the host respiratory tract and prevent IAV from spreading through the suppression of airway proteases at the level of the nasal mucosa. Thus, intranasal delivery of the human nasal commensal *S. epidermidis* represents a potential therapeutic approach for treating respiratory viral infections via the change of cellular environment in respiratory tract.

## Material and method

### Ethics statement

Participation was voluntary, with written informed consent obtained from all subjects. The Institutional Review Board of the Seoul National University College of Medicine approved the protocol for this study (IRB #C2012248 [943]). *In vivo* experiments with C57BL/6J mice were carried out according to guidelines approved by the Institutional Review Board of the Seoul National University College of Medicine (IACUC #2016-0093).

### Nasal mucus microbiome characterization

The mucus from the middle turbinate of healthy volunteers was collected individually using sterile 3M Quick swabs (3M Microbiology Products, ST Paul, MN, USA) from 4 subjects using a rigid 0-degree endoscope in an operating room. The swabs with mucus were fixed in a fixative solution and were transported immediately to the laboratory for identification and subsequent microbial analysis. For bacterial colony isolation, the mucus was placed on lysogeny broth (LB) plates. After 2 days incubation, bacterial colonies were obtained from the LB plates and the species of each colony were identified using GS-FLX 454 pyrosequencing by 16S rRNA gene amplification (17). *S. epidermidis* strains (N1-N4) were isolated from four individuals and 4 strains were used in the study.

### Viruses and reagents

*Influenza A virus* strain A/Wilson-Smith/1933 H1N1 (IAV A/WS/33; ATCC, Manassas, VA, USA) was used in this study to induce acute viral lung infection. Virus stocks were grown in Madin-Darby canine kidney cells in virus growth medium according to a standard procedure (3).

### Cell culture

Normal human nasal epithelial (NHNE) cells were cultured as described previously (55).

### Real-time PCR

NHNE cells were infected with WS/33 (H1N1) for 2, 4, 8, 24 hr and total RNA was isolated using TRIzol (Life technology, Seoul, Korea). cDNA was synthesized from 3 μg of RNA with random hexamer primers and Moloney murine leukemia virus reverse transcriptase (Perkin Elmer Life Sciences, Waltham, MA, USA and Roche Applied Science, Indianapolis, IN, USA).

### Quantification of secreted proteins

Secreted human urokinase plasminogen activator and Serpine1 were quantified using human urokinase plasminogen activator (DY1310) and human Serpine1 (DY1786) DuoSet ELISA kits (R&D Systems, Minneapolis, MN USA), respectively. The working range of the assays was 62.5-4000 pg/ml.

### Viral titer determination

Viral titers were determined using a plaque assay. Virus samples were serially diluted with PBS. Confluent monolayers of MDCK cells in six-well plates were washed twice with PBS and then infected in duplicate with 250 μl/well of each virus dilution.

### Western blot analysis

The protein levels of IAV NP, HAT, and Serpine1 were assessed using western blot analysis. The NHNE cells and were lysed with 2X lysis buffer (250mM Tris-Cl (pH6.5), 2% SDS, 4% β-mercaptoethanol, 0.02% bromophenol blue, and 10% glycerol).

### Murine infection model

Male C57BL/6J (B6) mice (Orientalbio, Seoul, Korea) aged 7 weeks (19–23 g) were used as wild-type (WT) mice.

### Immunohistochemistry and histologic analysis

Lung tissue was fixed in 10% (vol/vol) neutral buffered formalin and embedded in paraffin. Paraffin-embedded tissue slices were stained with hematoxylin/eosin (H&E) or periodic acid–Schiff (PAS) solution (Sigma, Deisenhofen, Germany). Histopathologic analysis of inflammatory cells in H&E-stained lung sections was performed in a blinded fashion using a semi-quantitative scoring system as previously described (56).

### LC-MS/MS

All LC-MS/MS analyses (DDA and DIA) were performed with the aid of a Quadrupole Orbitrap mass spectrometer (Q-Exactive plus; Thermo Fisher Scientific) coupled to an Ultimate 3000 RSLC system (Dionex, Sunnyvale, CA, USA) via a nanoelectrospray source, according to a modified version of the procedure described earlier (57).

### Data processing for label-free quantification

Mass spectra were processed with the aid of MaxQuant software (version 1.5.3.1). MS/MS spectra were analyzed with the aid of the Andromeda search engine (58).

### DIA MS data processing

To generate spectral libraries, we performed 12 urine DDA measurements and compared the spectra with those in the Maxquant Uniprot Human Database and the iRT standard peptide sequences. The spectral library (derived using individual the DIA data) was generated with the aid of Spectronaut ver. 10 software (Biognosys, Schlieren, Switzerland).

### Statistical analyses

For in vitro study, at least three independent experiments were performed with cultured cells from each donor, and the results are presented as the mean value ± standard deviation (SD) of triplicate cultures. Differences between treatment groups were evaluated by analysis of variance (ANOVA) with a *post hoc* test. We present the *in vivo* results of real-time PCR, plaque assays, and ELISA as mean values ± SD from five individual mice. Statistical analyses were performed with GraphPad Prism software (version 5; GraphPad Software, La Jolla, CA, USA). A *p*-value <0.05 was considered to be statistically significant.

## ACKNOWLEDGEMENT

H.J.K., and A.J., conceived the study and designed the experiments. J.W., C.H.C., and H.J.K. carried out the study including sample collection and sample preparation. K.M.L., and S.S.Y. performed additional work, design and data analysis. H.J.K., S.S.Y., and J.Y.C. drafted the manuscript.

